# The Kinship Formula: inferring the numbers of all kin from any structured population projection model

**DOI:** 10.1101/2023.03.29.534757

**Authors:** Christophe F. D. Coste

**Affiliations:** Biosciences Department, University of Swansea, SA1 8EN, UK; Centre for Biodiversity Dynamics, Department of Biology, NTNU, NO-7491, Trondheim, Norway; Unité Eco-anthropologie (EA), Muséum National d’Histoire Naturelle, CNRS, Université Paris Diderot, F-75016, Paris, France

**Author notes:** **Corresponding author:** Christophe F.D. Coste, Department of Biosciences, Swansea University, Singleton Park Campus Swansea, SA2 8PP, UK.

## Abstract

Structured population projection models are fundamental to many fields of science. They enable abundance forecasting for populations categorized by various traits such as age (for demography), patch (for spatial ecology), genotype (for genetics), infectious stage (for epidemiology) or capital (economics). The demography of a structured population, determined by the transition rates (e.g., survival, fertility) between its various states, also shapes its relatedness – or kinship – structure. This structure (a probabilistic genealogy) is crucial for understanding how individuals are related to the rest of the population and affects effective population size, inclusive fitness, inbreeding, pedigrees, relatedness, familial structures, etc. Despite its significance, the relationship between demography and kinship remains under-explored. By incorporating the generation number as a trait into the population structure, we derive the Kinship Formula, yielding the expected number of any kin for any structured population. This formula is simple to implement and fast to compute, even for complex models. Most importantly, it promises significant theoretical advances. From the Kinship Formula, one can, for instance, assess the impact of embedded processes (e.g., dispersal, inheritance, growth) on kinship, compute mean population relatedness and the eventual number of kin (including kin already dead or not born yet). The Kinship Formula derived here stems from a one-sex constant environment framework. Its simplicity should allow for extensions to include environmental and demographic stochasticity as well as two-sex models.

**Data accessibility statement:** No new data are used. All data used to illustrate the method are public. The code related to the main text (R and Matlab) can be found in Supplementary Materials (S.M.VIII)

## Introduction

Structured population projection models are crucial for many scientific fields. They enable the projection, over time, of populations categorised by one or several traits. This can be, for instance, a population of humans structured by age, sparrows by stage and body mass, susceptible individuals – in an epidemic – structured by infectious status, households by age and capital, drosophilas by genotype, cells by cancerous status, meta-populations by geographical patches. From the transition rates between all pair of states, one can predict population dynamics, future trait distribution, extinction probability of the population, and more. Such analyses are the bread and butter of ecology, epidemiology, genetics, anthropology, micro-economics and certain areas of physics.

The structured demography of a population not only informs about its future but also sheds light on its past history and current kinship (or relatedness) structure. The number of kin (or collaterals) an individual has is determined by its own reproduction, as well as the survival and reproduction of its ancestors and their descendants (1). Understanding the influence of population dynamics on relatedness structure is crucial across many fields and central to the emerging field of Kinship Demography (2). For example, in genetics, kinship structure affects inbreeding levels and effective population size (3; 4). In behavioral ecology, it influences how individuals may compete or cooperate and it determines their inclusive fitness (5; 6; 7). In anthropology and micro-economics, it combines with cultural norms to define familial structures (8; 9; 10). In epidemiology, the “kin” – individuals infected by a common source – can form clusters which are crucial for epidemic predictions (see, e.g., 11; 12; 13; 14). In population ecology and demography, the ability to predict kinship structures becomes increasingly crucial with the rising availability of pedigrees and genealogies (see, e.g., 15; 16; 17).

For simple, unstructured models with non-overlapping generations, population geneticists have developed tools to study the distributions of ancestors and collaterals (18; 19; 20; 21; 22). For age-structured, generation-overlapping populations, Le Bras published a method to trace probabilistic ancestry (23). Goodman, Keyfitz and Pullum extended this work (24) to provide the expected number of a few kin (ancestors *and* collaterals: sisters, aunts, mother, grand-mother, daughters and cousins) in a landmark study (2; 25). However this method – together with its matrix form (26) – focuses solely on age-structured populations and treats each kin type separately, which reduces its generalizability. Researchers interested in populations structured by traits other than age have therefore often resorted to simulations (or micro-simulations, via, e.g., the SocSim software, see 27; 28). Because of these limitations, this approach hinders Kinship Demography from fully exploring the impacts of specific processes (demographic, ecological, cultural, genetic) on relatedness, inbreeding or familial structure. It does not allow, for instance, to understand how dispersal, growth or inheritance affect the kinship structure of a population. To address this gap, Coste and collaborators recently published a closed-form formula for predicting the number of any kin within any structured population (29). While this formula represents significant progress for Kinship Demography, it lacks the simplicity and generalizability needed to be the theoretical bridge between demography and kinship that has been called for by demographers (24), mathematicians (30) and evolutionary ecologists (2). Although more general than the aforementioned alternative approaches, it remains bulky and complex to analyse theoretically.

Here, we introduce a model that structures the population both by the original traits and by generation. This leads to a substantial simplification of Coste et al’s formulation (29). By leveraging the genealogical Markov chains – developed by Demetrius (31; 32) and studied by Tuljapurkar (33; 34) – and the generation structure, we derive a simple formula (the Kinship Formula, eq.3) for the expected number of any kin from any structured population projection model. The Kinship Formula applies to close but also (very) distant kin and provides the structure of the kin network (e.g., their spatial dispersion or age distribution). Unlike other approaches, it enables the exact computation of the expected number of kin for a stage-structured population, which we illustrate for a ground squirrel population. The Kinship Formula facilitates rapid computation (even for a very complex model) and should prove crucial for sound extensions to more intricate frameworks (e.g., stochasticity, two-sex models). We demonstrate its utility by calculating the eventual number of kin. Further we show how the unstructured kinship (the number of kin irrespective of their state), when compared with the kinship of the unstructured population, enables analysis of the effects of traits on kinship. Finally we provide a link between the individual measure of the number of kin and the population measure of relatedness.

### The Framework

Consider a general, structured and generation-overlapping, population. The structure may consist of one or several traits (see S.M.I, for the concept of traits in projection models). We limit ourselves to one-sex models here and to processes that are constant in time. To be as general as possible, we consider a discrete structure and a discrete time-step (see S.M.II). The discrete structure consists of *s* classes (or states). This population can be projected over time, via 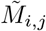, the stochastic process yielding the number of individuals in class *i* stemming from an individual in class *j*, over the chosen time-step. Process 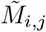 can be decomposed into a survival (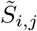 “producing” the same individual) and a fertility ( 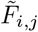, producing new individuals) components: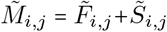. The deterministic expectations of this process – we denote 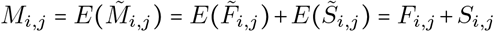 – projects the deterministic population vector ***n*** over time: *n*_*i*_(*t +* 1)= ∑_*j*_ *M*_*i*,*j*_*n*_*j*_(*t*) which, in matrix form, is ***n (****t+* 1)=**M*n*(***t*)=(**F***+* **S)*n*(***t*). Matrix **M** is the so-called population projection matrix (35), which elements, the *M*_*i*,*j*_ are called the transition rates. We consider, without any loss of generality, that the processes of reproduction and survival are independent and identically distributed (see S.M.III). Notation-wise, stochastic processes are noted with a tilde, and expectations without, vectors (lower case) and matrices (upper case) are in bold while scalars are in plain text. The population projection matrices **S** and **F** are the building blocks of the Kinship Formula (eq.3) inferring the expected number of kin (structured by kin type and by class) of focal individual *ego* (categorized by its class). They will allow us, in a generation-structured population (eqs.1 and 2) which makes it possible to consider all kin of *ego* simultaneously, to go back in time towards the distribution of the ancestors of *ego* via their genealogical Markov chains, and from there to count the descendants of these ancestors.

#### Genealogical Markov Chains

At the demographic stable state, the population growth rate and the vector of relative abundances (denoted respectively *λ* and ***w***) correspond to the maximum eigenvalue of **M** and its associated right-eigenvector (scaled to sum to 1): **M*w***=*λ****w***. While **M** projects the population forward in time, its genealogical Markov chain (31; 32) projects it backwards in time. The transition matrix of the genealogical Markov chain associated with **M** is **P**, where 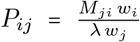 is the probability that the ancestor (via survival or reproduction) of an individual in class *j* was in class *i* at the previous time-step (see appendix 1). Similarly to the projection matrix **M**, the transitions of **P** can be decomposed according to survival and fertility: **P** = **P**_**S**_ + **P**_**F**_ (eq.13, in app.1).

#### Same-litter daughters and sisters

When projecting individuals forward in time, we must consider the number of “same-litter sisters” – the sisters born at the same time than *ego* – which depends on the stochastic fertility process 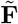 (and not only deterministic **F**), a fact already accounted for by Goodman, Keyfitz and Pullum (24). Another component of the Kinship Formula is therefore the number of “same-litter-daughters” 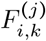, the expected number of supplemental offspring (in class *i*) produced by a mother (in class *k*) knowing she has produced one (in class *j*) (eq.14 in app.2). In an operation, that mimics, at the level of one generation, the general pattern yielding the Kinship Formula, we generate, from **P**_**F**_ and 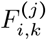, the same-litter sister matrix **Z**. Its *Z*_*i*,*j*_ element is the number of “same-litter sisters” in class *j* of *ego* in class *i* (29) and corresponds to 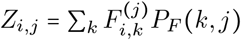 (see eq.15). In many cases, in practice, the number of same-litter daughters does not depend on the class of the individual known to have been produced (see below and S.M.IV), which yields a simpler Kinship Formula (eq.8).

#### Generation structured population

We now incorporate an additional trait into the population structure: the *generation number*. We consider a population vector that is 2-dimensional, that is, made of two traits: the first trait corresponds to the *s* classes of the original structure (which can itself correspond to several traits that we consider combined into one) and the second trait to the *generation number* (see S.M.I). We turn this second dimension, which is in principle relative and infinite, into an absolute and finite metric by setting the generation number of the *most recent common ancestor* of the kin and *ego* to 1 and by providing trait *generation number* with an arbitrary number (*g*_*m*_) of classes. The multitrait population vector (or tensor) needs to be vectorised (to be made one-dimensional) in order to be projected over time: individuals in class *c*_*i*_ and generation *g*_*i*_ are to be found in the *i*^*th*^ entry of 𝕟, the new population vector, where *i* = *c*_*i*_+*s*(*g*_*i*_−1); we will denote this entry equivalently 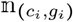 or 𝕟_*i*_. We denote 𝔽 and 𝕊 the generation-structured matrices projecting 𝕟 forward in time and ℙ_𝔽_ and ℙ_𝕊_ the generation-structured projectors backward in time. Such multitrait population projection matrices take the appearance of block-matrices; here, they are of size *s*.*g*_*m*_×*s*.*g*_*m*_ made of *g*_*m*_×*g*_*m*_ blocks of size *s*×*s*. When the trait added on top of an existing structure does not affect the demographic processes, as for trait *generation number* here, the new model can be built simply using the tensor (or Kronecker) product ⊗ . Here the generation number is constant via survival and increases by 1 via reproduction, and thus

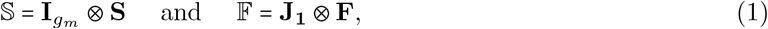

with **J**_**1**_ representing a zero matrix of size *g*_*m*_×*g*_*m*_ except for ones on the sub-diagonal and 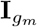 the identity matrix of size *g*_*m*_. Block-matrix 𝕄=𝔽+𝕊 has therefore **S** as its diagonal blocks and **F** as its sub-diagonal blocks; for *g*_*m*_ = 3, this is:

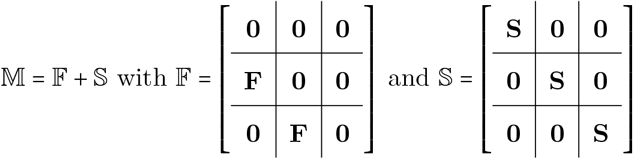

Similarly, we have

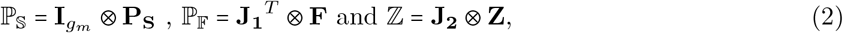

with **J**_**2**_ the zero *g*_*m*_ × *g*_*m*_ zero-matrix but for a 1 in the (2,2) position.

#### Characterisation of kin relationships and the Kinship Matrix

We aim to derive a mathematical object, we call the Kinship Matrix 𝕂, that, like 𝔽 or ℙ_𝕊_, is generation-structured and which 𝕂((*c*_*i*_, *g*_*i*_), (*c*_*j*_, *g*_*j*_)) entry represents the expected number of (*c*_*j*_, *g*_*j*_) descendants of the ancestor (in generation 1) of *ego* in state (*c*_*j*_, *g*_*j*_). These are referred to as the (*g*_*i*_, *g*_*j*_) -kin (in class *c*_*i*_) of *ego* (in class *c*_*j*_). This approach provides a complete characterisation of all possible kin relationships: the (*g*_*i*_, *g*_*j*_) -kin are the *g*_*i*_-descendants of the *g*_*j*_-ancestor of *ego*, where the *g*_*i*_-descendants lie *g*_*i*_ − 1 generations down from the ancestor (e.g., the 3-descendants are the grand-daughters) and the *g*_*j*_-ancestor is to be found *g*_*j*_ − 1 generations up from *ego* (e.g., the 2-ancestor is the mother). Thus, the (1, 2)-kin is the mother, the (3, 1)-kin the grand-daughters and the (3, 3)-kin the (first) cousins. We illustrate this characterisation of kin relationships in fig.1, which corresponds to a figure used for the same purpose in (36; 37; 38) but with an indexation that starts at (1, 1) for *ego* (instead of (0, 0) in these papers). This characterisation corresponds to the various blocks of the Kinship Matrix: the (*g*_*i*_, *g*_*j*_) -block of 𝕂 corresponds to the number of (*g*_*i*_, *g*_*j*_)-kin. In particular, the first block-row of 𝕂 corresponds to the ancestors of *ego* (including itself), and each related block-column corresponds to the descendants of that ancestor (see eq.9 for a numerical illustration).

**Figure 1.**
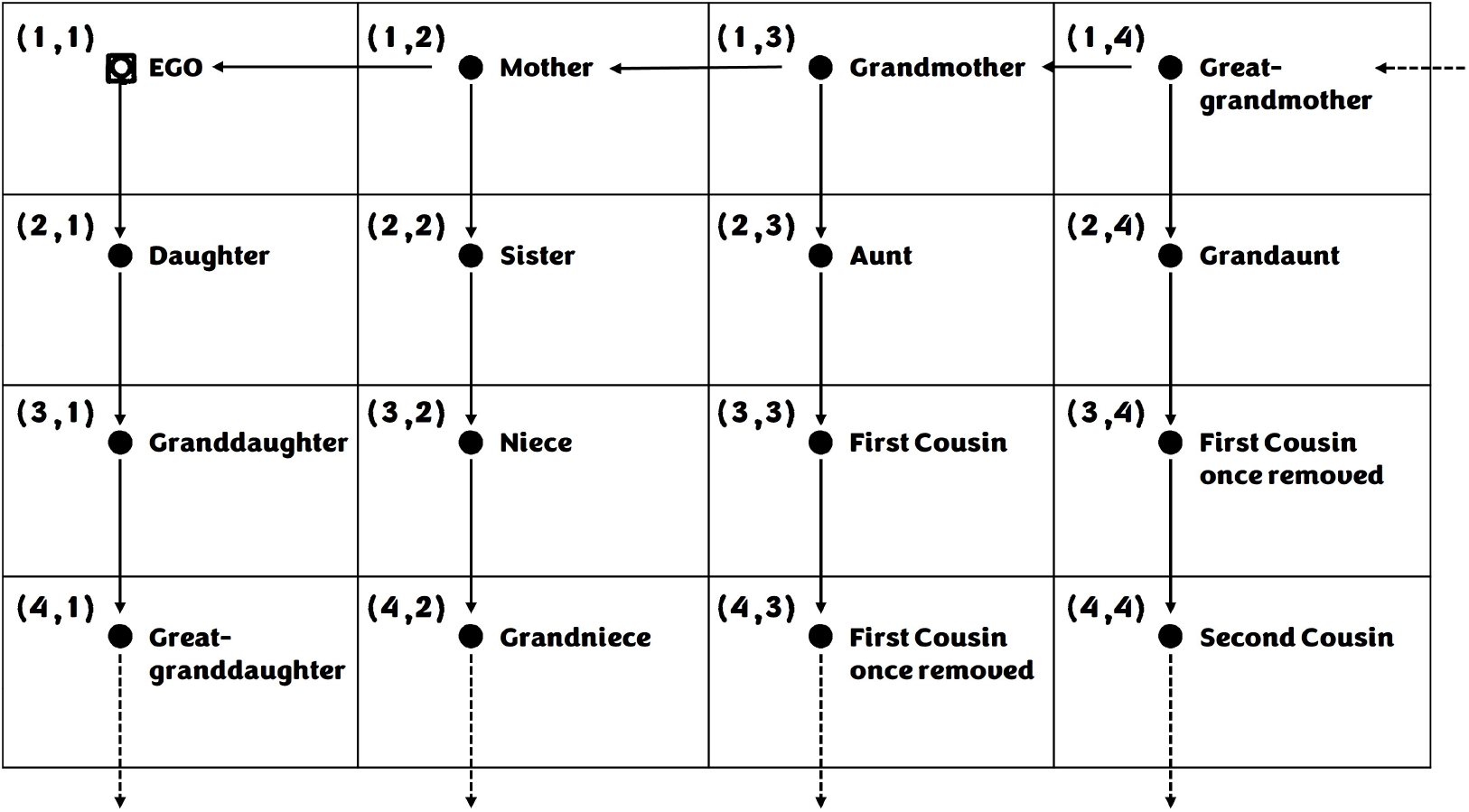
Characterisation of (*g*_*i*_, *g*_*j*_)-kin relationships, adapted from (36).

## Results

### The Kinship Formula

#### Theorem 1.

If **S** converges (i.e., if lim_*t*→+∞_ **S**^*t*^ = **0**), the Kinship Matrix 𝕂 is the unique solution of

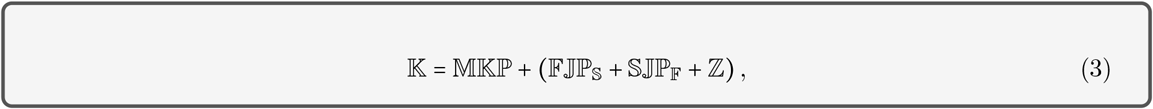

where 𝕁=**J**_**3**_⊗**I**_*s*_ is the zero *sg*_*m*_×*sg*_*m*_ block-matrix but for the diagonal of the first block made of 1s and **J**_**3**_, the zero *g*_*m*_×*g*_*m*_ matrix with 1 in the (1,1) position.

We demonstrate, below, that 𝕂 is indeed a solution of eq.3 and, in appendix 5, we establish the existence and uniqueness of this solution, provided that **S** converges. This fundamental condition for most projection models implies that every individual dies eventually. The formulation of eq.3 does not allow for a direct computation of 𝕂 (it can be found on both sides of the equation) which we provide in the next section (eqs.5,6).

#### proof and interpretation

Equation 3 states that the kin of *ego* (the 𝕂 on the left-hand side) are the descendants of the kin of *ego*’s immediate ancestor (its ancestor one time-step ago), denoted as 𝔹=𝕄𝕂ℙ, plus additional kin denoted as 𝔸=(𝔽𝕁ℙ_𝕊_+𝕊𝕁ℙ_𝔽_+ℤ ).

Consider 𝕂((*c*_*i*_, *g*_*i*_),(*c*_*j*_, *g*_*j*_)) representing the number of (*g*_*i*_, *g*_*j*_)-kin in class *c*_*i*_ of *ego* in class *c*_*j*_. The immediate ancestor of *ego* is in class *c*_*a*_ and generation *g*_*a*_ with probability ℙ((*c*_*a*_, *g*_*a*_),*(c*_*j*_, *g*_*j*_) There are two cases: either *ego*’s immediate ancestor is *ego* itself (if alive at the previous time-step) – then *g*_*a*_ = *g*_*j*_ and ℙ((*c*_*a*_, *g*_*a*_),(*c*_*j*_, *g*_*j*)_)= *P*_*s*_*(c*_*a*_, *c*_*j*_)– or *ego* has just been born and *ego*’s immediate ancestor is *ego*’s mother – then *g*_*a*_*=g*_*j*_ *−*1 and ℙ((*c*_*a*_, *g*_*a*_),*(c*_*j*_, *g*_*j*_*)=P*_*f*_*(c*_*a*_, *c*_*j*_). Note that if *g*_*j*_=1, we do not consider *ego*’s mother’s kin as the (*g*_*j*_, 1)-kin of *ego* are its direct descendants. At the stable state, the number of (*g*_*b*_, *g*_*a*_)-kin in class *c*_*b*_ of the immediate ancestor (in class *c*_*a*_) is 𝕂 ((*c*_*b*_, *g*_*b*_),(*c*_*a*_, *g*_*a*_)). We project the kin of the immediate ancestor towards the present day via 𝕄: the expected number of (*g*_*i*_, *g*_*j*_)-kin in class *c*_*i*_ for *ego* in class *c*_*j*_ corresponding to the immediate descendants of its immediate ancestor is 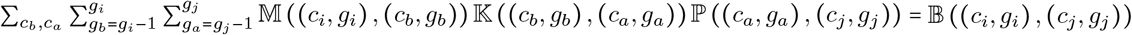, If *g*_*a*_*>*1, that is, if *ego*’s immediate ancestor is not the most recent common ancestor of *ego* and the kin, then this counts all (*g*_*i*_, *g*_*j*)_ -kin of *ego* today and 𝕂((*c*_*i*_, *g*_*i*_),(*c*_*j*_, *g*_*j*_)) = 𝔹((*c*_*i*_, *g*_*i*_, *c*_*j*_, *g*_*j*_)). If *g*_*a*_*=*1 and *ego*’s ancestor is *ego* itself (*g*_*j*_*=*1) then 𝔹 counts all kin of *ego* today but for those that *ego* has just produced which correspond to 𝔽𝕁ℙ_𝕊_. If *g*_*a*_ = 1 and *ego*’s ancestor is *ego*’s mother (*g*_*j*_*=*2) then 𝔹 counts all kin of *ego* today but for those that *ego*’s mother has just produced (*ego*’s same-litter sisters) and *ego*’s mother herself (*ego* is not a kin of *ego*); the sum of these two components corresponds to ℤ+𝕊𝕁ℙ_𝔽_. Overall, we account for these aditional kin via 𝔸=(𝔽𝕁ℙ_𝕊_+𝕊𝕁ℙ_𝔽_ +ℤ), where dummy matrix 𝕁 ensures that the number of additional kin is non-zero only for *g*_*a*_*=*1. Therefore, ∀*c*_*i*_, *c*_*j*_, *g*_*i*_, *g*_*j*_, 𝕂 ((*c*_*i*_, *g*_*i*_), (*c*_*j*_, *g*_*j*_)) = (𝔸 + 𝔹) ((*c*_*i*_, *g*_*i*_), (*c*_*j*_, *g*_*j*_)), which completes the proof.

The three-way decomposition of 𝔸 corresponds to that of kin via older, younger and same-litter sisters implemented already in the foundational works of Kinship Demography (24; 37; 29). Distinguishing between older and younger sisters is essential because the knowledge of the ancestor’s status varies when considering potential siblings born before (the ancestor is known to be alive) of after (the ancestor may not be alive) *ego*’s birth. Additionally, the inclusion of the same-litter daughters is necessary due to their aforementioned dependence on the stochastic reproductive process (29). This formulation of the Kinship Formula (eq.3) lays the groundwork for extending the framework to incorporate individual stochasticity and compute the variance and other moments of the number of kin.

### Computation of the Kinship Matrix and alternative formulations

We will now present an alternative expression of the Kinship Formula enabling direct computation of 𝕂. First, we simplify eq.3 by substituting the components of **P** by their expressions as functions of **M**’s components (see eq.13 in app.1). Let **D**_***w***_ be the diagonal matrix with ***w*** on its diagonal and 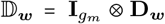. Letting 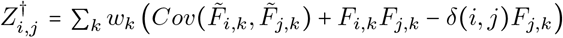 (eq.16 in app.3), ℤ^†^ = **J**_**2**_ ⊗ **Z**^†^,and 𝕂_***w***_ = 𝕂𝔻_***w***_, we arrive at an equivalent formulation of the Kinship Formula where the transition matrix of the genealogical Markov chain is not explicit (see appendix 3 for details):

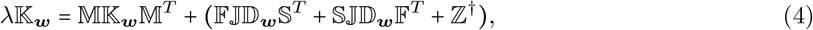

While eq.4 may seem less intuitive than its counterpart eq.3 due to the absence of the genealogical Markov chain, it offers greater analytical tractability. This expression should facilitate the computation of the sensitivity of the kinship structure to the demographic rates and the integration of varying environments.Notably, when expressed as 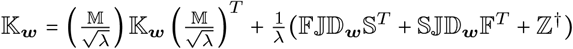, equation 4 becomes a Lyapunov (or Stein) equation (39), extensively studied in signal processing and control theory (40). The use of vectorization to solve Lyapunov equations is a classic technique (41) and we will now proceed with this operation.

#### Vectorization of the Kinship Formula

Let us now introduce the ***vec*** operator which turns a matrix into a vector by “stacking” its columns into a column vector (42). From eq.4, this operator allows to provide an alternative and equivalent version of the Kinship Formula (see apps.4,5):

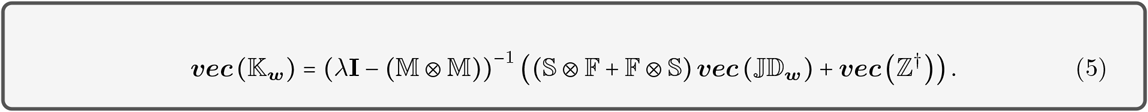

From this, we can readily obtain the Kinship Matrix, using the inverse of the ***vec*** operator, denoted 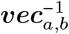, which turns a vector ***d*** of size *ab* into an *a* × *b* matrix such that 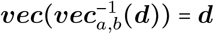 :

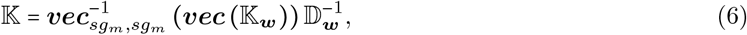

which is simple to implement and fast to compute. We illustrate this (below and in S.M.V) and provide the code to compute 𝕂 from eq.6, for both Matlab and R, in S.M.VIII.

#### The Kinship formula as an infinite sum

The Kinship Formula can also be equivalently expressed as (see S.M.VI)

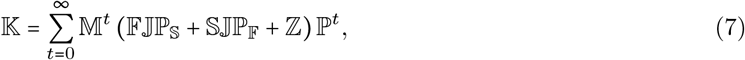

Computationally, this formulation can only provide the exact number of kin when the sum is finite, i.e., when a maximum age *ω* is embedded in the projection matrix (such that **S**^*ω*^ = **0**). In this case 𝕂 simplifies to: 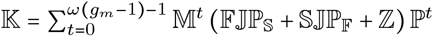. While this equation is of lesser interest than its alternatives, it remains useful. For instance, we employ this this approach when computing the eventual number of kin (see below and S.M.VII).

### Particular Reproductive Cases

In some particular cases, that are likely to constitute the majority of cases in practice, the number of same-litter-daughters 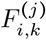 is independent from the class (*j*) of the offspring known to have been produced. This simplifies computation, allowing the number of same-litter-daughters to be represented by a matrix **F**^∗^. In its generation-structure formulation, 𝔽^∗^ = **J**_**1**_⊗ **F**^∗^, it allows computing directly ℤ=𝔽^∗^𝕁ℙ_𝔽_ and ℤ^†^ = 𝔽^∗^𝕁𝔻_***w***_𝔽^*T*^, which can then be incorporated into the various versions of the Kinship Formula. For eq. 3, this is 𝕂 = 𝕄𝕂ℙ + (𝔽𝕁ℙ_𝕊_ + 𝕊𝕁ℙ_𝔽_ + 𝔽^∗^𝕁ℙ_𝔽_) and for eq.5:

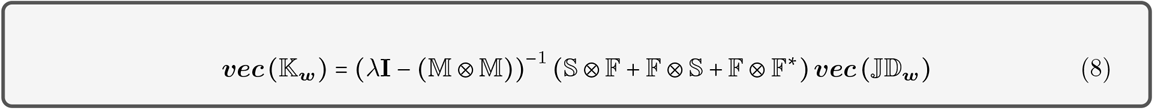

We now consider three such cases (see S.M.IV). These are

#### Bernoulli Reproduction

Individuals produce at most 1 offspring per time-step, hence **F**^∗^ = **0**.

#### Independent Poisson reproduction

Here, the number and class of supplemental offspring is independent from the number and classes of offspring already produced and therefore **F**^∗^ = **F**.

**One offspring class** (say, class 1) then, from eq.14, **F**^∗^ is such that 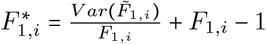.

### Illustration

In S.M.V, we illustrate the computation of the number of kin for a population of ground squirrels, *Spermophilus dauricus*, structured by 3 stages. From the life table in (43) and for *g*_*m*_=3, we obtain the following Kinship Matrix (via eqs.8 and 6):

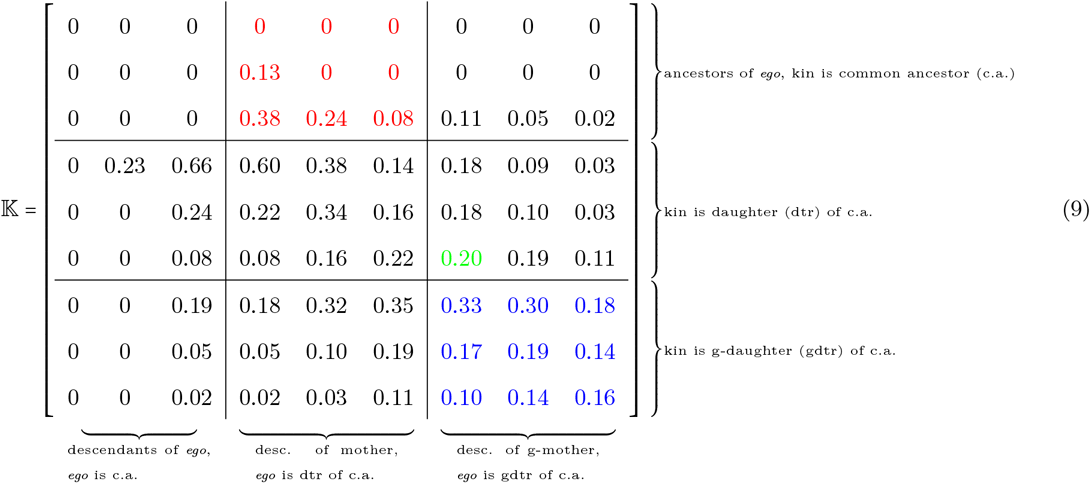

In the (1, 2) block of eq.9, highlighted in red, we find the expected number of mothers, i.e., the probability for *ego* to have its mother alive (see fig.1). The Kinship Matrix 𝕂 indicates, for instance, that a yearling (an individual in class 1) has a 13% probability for its mother to be alive *and* in class 2. In the (3, 3) block, highlighted in blue, we observe the expected number of cousins: a class 2 individual, on average, has 0.19 cousins in class 2. The element in green, corresponds to the ((3, 2), (1, 3)) entry of 𝕂, representing the expected number of (2, 3)-kin, or aunts, in class 3 for *ego* in class 1. Notably, the kin numbers in eq.9 are exact, contrasting with the approximation in (29).

### Extensions

#### Unstructured Kinship

The Kinship Matrix 𝕂 is structured by class and type of kin. It is also possible to consider the unstructured kinship of this structured population, meaning the number of kin regardless of their class for an *ego* chosen at random from the entire population. Let 𝕂^*u*^(*g*_*i*_, *g*_*j*_) be the expected number of (*g*_*i*_, *g*_*j*_)-kin of such an individual. At the stable state, the probability for *ego* to be in class *c*_*j*_ is 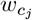 and the number of its *g*_*i*_ kin is the sum over all *c*_*i*_, therefore 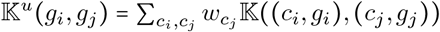, or, in matrix form:

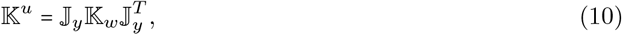

with 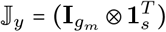 the *g*_*m*_ × *sg*_*m*_ matrix where the *i g*_*m*_ × *s* block is zero but for the *i*^th^ row filled with ones, **1**_*s*_ the vertical vector (of size *s*) of ones and ^*T*^ denoting matrix transposition.

#### The Unstructured Kinship Matrix 𝕂^*u*^ is symmetrical

This is a key result from this manuscript:

##### Theorem 2.

For *ego* drawn at random in the population at the stable state, its expected number of (*g*_*i*_, *g*_*j*_ )-kin is equal to its expected number of (*g*_*j*_, *g*_*i*_)-kin: ∀*g*_*i*_, *g*_*j*_, 𝕂^*u*^(*g*_*i*_, *g*_*j*_ ) = 𝕂^*u*^(*g*_*j*_, *g*_*i*_) or, in matrix form:

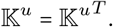

In other words, the probability for an individual to have its mother alive is equal to the expected number of its daughters, and the expected number of nieces is equal to that of aunts. This result is intuitive: a (*g*_*i*_, *g*_*j*_) kin relationship for *ego* corresponds to a (*g*_*j*_, *g*_*i*_) kin relationship for the kin. In a population with *n* individuals, among the *n(n −*1)(directional) relationship pairs, there will be as many (*g*_*i*_, *g*_*j*_ ) kin relationships as (*g*_*j*_, *g*_*i*_) ones: 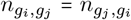. At the individual level, this implies that 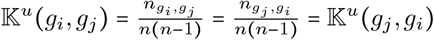

**proof** The symmetry is absent from 𝕂 itself, but present in 𝕂_*w*_. Indeed, as 𝔻_***w***_𝕁 = 𝕁𝔻_***w***_, we have (𝔽𝕁𝔻_*w*_𝕊^*T*^)^*T*^ = 𝕊𝕁𝔻_*w*_𝔽^*T*^, From eq.16, it is clear that **Z**^†^ and therefore ℤ^†^ are symmetrical. From eq.4, this implies that *λ*𝕂_***w***_ − 𝕄𝕂_***w***_𝕄^*T*^ is symmetrical which leads to 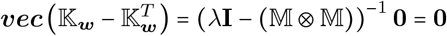 and therefore 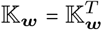. From eq.10, this further implies that 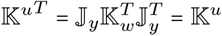, which completes the proof.

**illustration** For *g*_*m*_ = 3, the Unstructured Kinship Matrix for the ground squirrels population is (S.M.V):

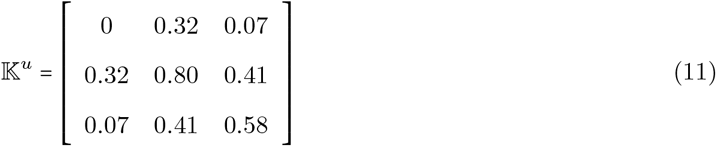

From eq.11, we can infer that an individual taken at random in the population, is expected to have= 0.80 sisters, 0.58 cousins, 0.41 nieces and aunts and 0.07 grand-daughters and grandmothers (i.e., the probability that its grandmother is alive is 0.07). We observe that 𝕂^*u*^ is indeed symmetrical. We can explore the number of kin at much larger *g*_*m*_ distances. In fig. 2, we show the expected (unstructured) number of kin for the population of ground squirrels (a declining population, *λ* < 1) for *g*_*m*_ = 300. Each curve corresponds to the number of (*g*_*i*_, *g*_*j*_) kin for a given *g*_*j*_, that is, to the descendants of the *g*_*j*_-ancestor (see fig.1). For *g*_*j*_*=*1 (in dark blue), it corresponds to the numbers of the various direct *g*_*i*_-descendants of *ego*, which (from th.2) is also the number of its direct *g*_*i*_-ancestors. Dots correspond to the elements already appearing in 𝕂^*u*^ for *g*_*m*_ = 3 of eq.11. After a “transient” period corresponding to close kin (low *g*_*i*_ and *g*_*j*_), we observe that, for low or high *g*_*i*_, *ego* has no (*g*_*i*_, *g*_*j*_) kin alive (they are are already dead or not born yet) and that 𝕂^*u*^ (*g*_*i*_, *g*_*j*_), for a given *g*_*j*_, reaches a maximum at *g*_*i*_*=g*_*j*_, corresponding to *ego*’s 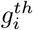 cousins (we point to an asymptotic analysis of kinship for unstructured populations in (29)).

**Figure 2.**
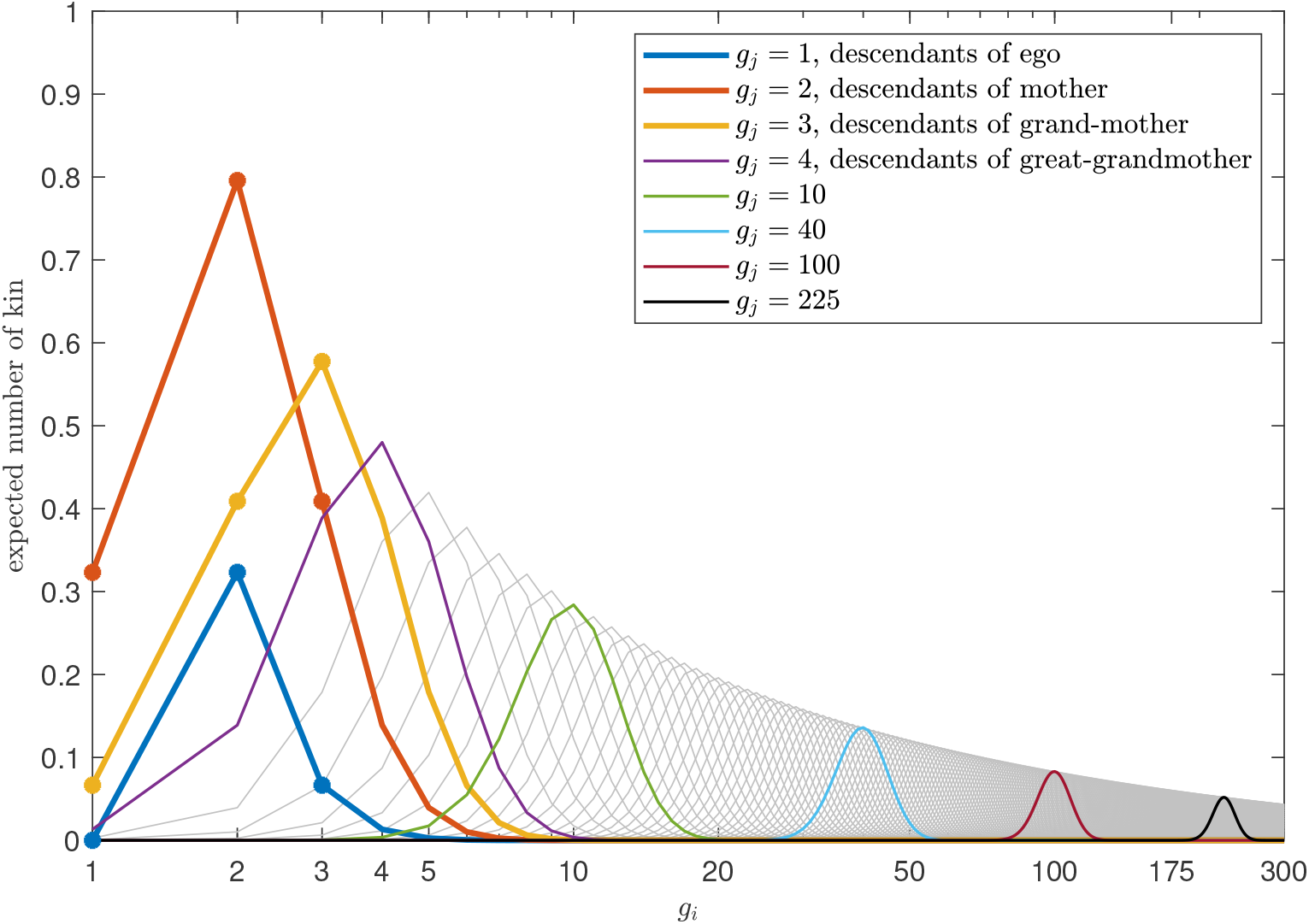
(unstructured) number of -kin, (*g*, *g* ) 𝕂^*u*^ (*g*_*i*_, *g*_*j*_),, for the ground squirrel population (illustration) for 1≤ *g*_*i*_, *g*_*j*_ ≤ 300 (note: the x axis, *g*_*i*_, is on a *log* scale). Each curve represents the descendants of the *g*_*j*_-ancestor of *ego* for a given *g*_*j*_ (see fig.1). Dots correspond to elements appearing in eq.11.

#### Unstructured kinship vs kinship of unstructured population

The Kinship Matrix offers a novel avenue for application of the Trait Level Analysis developed in (44). This methodology assesses the sensitivity of a population model to a specific trait by comparing the model to its version without (or “folded on”) that trait (see S.M.I). Within this multitrait theoretical framework, 𝕂^*u*^ represents the (*class ×generation*)-matrix 𝕂, *folded* over trait *class*. By contrasting 𝕂^*u*^(**M**), the unstructured kinship of the structured model **M**, with the kinship of the unstructured population 𝕂 (**M**^**u**^)derived from **M** *folded* over trait *class* (denoted **M**^**u**^), we gain insights into how the process(es) embedded in the *class* heterogeneity influence the kinship structure. We illustrate this approach using the ground squirrel population in S.M.V, demonstrating that their survival senescence (together with the age-fertility pattern) leads to a reduction in the number of direct ascendants (mother, grand-mother) and descendants (daughters, grand- daughters) while simultaneously increasing the count of “indirect” kin, such as sisters, nieces, cousins and aunts.

#### Kinship and Relatedness

The Kinship Formula (eq.3) provides the number of kin in a one-sex population. Consequently, it is possible to relate the Kinship Matrix to the relatedness along the one-sex lineages of the population (45). This requires defining a “relatedness distance” *d* corresponding to the length of the path connecting two individuals (29). In a (*g*_*i*_, *g*_*j*_) kin relationship, the kin is (*g*_*i*_ −1) generations away from the most recent common ancestor and *ego* is (*g*_*j*_ −1) generations away from that ancestor. Therefore *d (g*_*i*_, *g*_*j*_) = (*g*_*i*_ −1)+(*g*_*j*_ −1)= *g*_*i*_*+g*_*j*_ −2. The coefficient of relatedness for a (*g*_*i*_, *g*_*j*_) kin relationship is given by 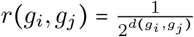 (46; 47), that is:

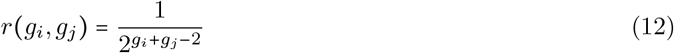

Each relatedness coefficient 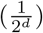 corresponds to the various entries on the (*d+*1)^*th*^ anti-diagonal of 𝕂^*u*^ (eq.10). For *d* = 2 (i.e., *g*_*i*_*+g*_*j*_*=* 4), it accounts for the grand-mother, the sisters and the grand-daughters and for *g*_*m*_ = 3 (as per eq11), it corresponds to the main anti-diagonal.

#### mean relatedness

These considerations extend the individual-centered kinship formula to population measures of relatedness. One common measure is the mean relatedness, which typically focuses on male or female sub-populations and considers the mean relatedness coefficient across all dyads (a dyad is a non-directional pair) for a given relatedness depth *g*_*m*_. First, consider 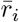, the *individual* mean relatedness (48). It can be obtained from *r*(*g*_*i*_, *g*_*j*_ ) and 𝕂^*u*^ via 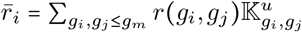, which, from eq.12, yields

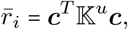

with vector ***c*** of length *g*_*m*_ such that 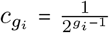 . From this individual relatedness, we can compute the population or *dyad* mean-relatedness 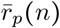 of a population of size n. There are *n*(*n* − 1) directional pairs of individual and therefore 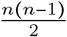 dyads in such a population. The total relatedness is 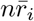, in which each dyad is counted twice. Hence, the population mean relatedness is

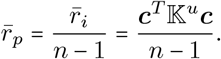

#### Eventual number of kin

The Kinship Formula provides the *current* number of kin of *ego*, where both *ego* and the kin are alive. In human demography and anthropology, where all kin matter including those not alive at the same time than *ego*, it is valuable to know the *eventual* number of kin: their total number encompassing those who died before *ego*’s birth and those who will be born after *ego*’s death. In S.M.VII, we show that this can be achieved by adding a class “death” to the original model structure. Applying this method to an unstructured, non-generation-overlapping population where individuals produce *R*_0_ offspring per generation and the expected number of sisters is *Z*, we obtain for *g*_*m*_*=*4, the following *eventual* number of kin:

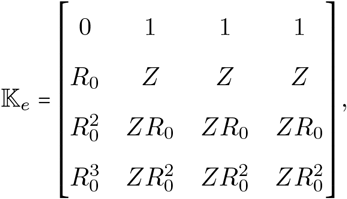

which is the same result obtained by Goodman et al.(24, fig 2) and by Pullum (37, p.550).

### Conclusion and Discussion

In this article, we present the Kinship Formula (eq.3), which calculates the expected number of kin, for any kin relationship and for any generation-overlapping structured population, in a one-sex constant environment framework. This formula utilizes the genealogical Markov chain (31), a model projecting populations backward in time, to produce the Kinship Matrix (the expected number of all kin of *ego* structured by kin type and the classes of the kin and *ego*), thereby extending results previously restricted to age-structured populations (24; 26) to any structured population. For example, the Kinship Matrix can analyze relatedness within spatially structured populations at both the patch and meta-population levels. Thanks to its generation structure, the Kinship Formula is closed-form and provides the exact number of all kin, up to an arbitrary relatedness distance, simultaneously. Both the genealogical Markov chain and the generation-structure are likely to have further importance for Kinship Demography in particular and evolutionary demography in general. Generation-structured multi-type branching processes will be central to studying the variance and distribution of the number of kin, extending the expectation analyzed here (manuscript in prep.). The genealogical Markov chain should prove important for extending results from age-structured models to any structured population, such as the Euler-Lotka equation, a crucial tools for evolutionary studies (49; 29).

We expect the simplicity of the Kinship Formula to appeal to population ecologists who wish to infer relatedness frequencies in their study population and compare them to available pedigrees. Demographers and anthropologists will benefit from its ability to compute the exact number of kin without resorting to simulations. For evolutionary and behavioral ecologists, the formula facilitates comparisons of relatedness structures across various social species, enabling analysis from an inclusive fitness perspective (50). Its simplicity allows for large-scale theoretical investigations of relatedness, as demonstrated in fig.2, which will be crucial for geneticists studying the distribution of identical-by-descent alleles and breeding values in a population (51; 52) In essence the Kinship Formula allows to extend to generation-overlapping structured populations the Kingman’s coalescent (20). Additionally, the Kinship Formula is valuable for micro- and family-economists, as it can quantify the effects of inheritance systems on the distribution of economic capital across extended families. Epidemiologists will find it useful for deriving information on the likelihood of an infected person “descending” from a “cluster” within a spatially-structured compartmental model. The Kinship Formula’s versatility arises from its applicability to any structured population, encompassing the two main types of individual heterogeneity: dynamic (classes are “stages”) and fixed (classes are “types”) heterogeneity (53; 54). It applies to (age- or) stage-structured models (55; 56), to genetic models where individuals are categorised by genotype or breeding value (51; 52), and to models combining both dynamic *and* fixed heterogeneity (57). As it allows to compare the unstructured relatedness to the relatedness of the unstructured population, the Kinship Formula is crucial for analyzing how different processes embedded within different traits affect the kinship network (44). For life historians, this is useful for understanding the consequences of trade-offs and the presence of various strategies on a population’s relatedness landscape.

The kinship formula provides the expected number of kin at the demographic stable state, for a one-sex population. Its simplicity makes it a foundational tool for developing extensions to more realistic frameworks, in a process that would mirror the historical progress in projection models: understanding the deterministic long-term growth rate as the maximum eigenvalue of a projection model (58), allowed to incorporate varying environments (59), individual stochasticity (60), two sexes (61) and compute its sensitivity (62). Ongoing and future works aim to follow the same path for Kinship Demography. The theoretical studies enabled by the Kinship Formula, such as investigating long-distance kinship structure (a study crucial for genetics, initiated in 29), hold significant potential across various fields interested in kinship structure and the behavior of collaterals in dynamical processes.

Finally, while these future extensions will enhance our understanding of how demography affects kinship composition, they represent only one aspect of Kinship Demography. This field aims to establish a comprehensive theory linking population dynamics and relatedness by examining both the effects of demography on kinship and the feedback of kinship on demography (2). In population ecology, the presence of kin significantly impacts individual fitness in many animal species, particularly in social species like whales and great apes (63; 64). In human societies, the effects of kin presence are notable, such as the extended care required for young (called altriciality, see 65) and the costs associated with caring for frail older relatives (66). Incorporating these effects into population demography has, for instance, reassessed the selection pressures on reproduction and survival in the context of menopause and susceptibility to late-onset diseases (68). The overarching goal of Kinship Demography is to generalize these ad-hoc studies to understand the feedback loop between demography and kinship, model populations via both kinship-dependent demography and demography-dependent kinship, and formulate a general theory of their co-evolution. These future investigations will rely heavily on the Kinship Formula and emulate the work of Chu on the co-evolution of inter-generational transfers (69), and the general approach used by May (70) and others when studying density-dependent population (where density, like kinship, is both an input and an output of the population model).

## Acknowledgments and funding sources

This study was supported by the Chair “Modélisation Mathématique et Biodiversité of Veolia – Ecole Polytechnique – MNHN–Fondation X”, by the Agence Nationale de la Recherche (grant ANR-18-CE02- 0011, MathKinD), by the Research Council of Norway (Centre of Excellence grant SFF-III, project 223257) and by the Natural Environment Research Council (grant no. NE/W006731/1). We thank Samuel Pavard and Mike Fowler for helpful comments and suggestions.

## Appendices

### 1 Genealogical Markov Chains

From *λ* and ***w***, the maximum eigenvalue of **M** and its associated right-eigenvector, the genealogical Markov chain of **M** (31; 32; 71; 34) allows to project the population backwards in time. We denote its transition matrix **P**, where 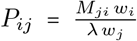 is the probability that the ancestor (via survival or reproduction) of an individual in class *j* was in class *i* at the previous time-step. Decomposed according to survival and fertility processes, this is **P** = **P**_**S**_ + **P**_**F**_, where

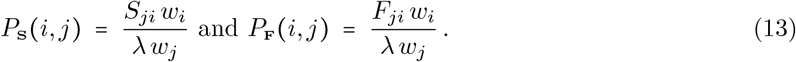

Note that, in order for backwards projection to obey the same rule of left-multiplication as projection matrices, we write **P** as a left stochastic matrix; **P** is therefore the transpose of the matrix of the same name in, e.g., (31) or (29).

#### Genealogical Markov chain and distribution of maternal age

The genealogical Markov chain generalises to any structure, the distribution of age of mothers (at birth-giving) used, for age-structured populations, in (24; 26). For a Leslie matrix (55; 72) – the projection matrix of a population structured by age – we get a **P**_**S**_ which is zero but for the sub-diagonal made of 1s (individuals older than the first age class necessary stem from their younger selves in the previous age class) and a **P**_**F**_, which is zero but for its first column which corresponds to the distribution of age of mothers.

### 2 Same-litter daughters and sisters

The number of same-litter-daughters 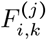 is the expected number of supplemental offspring produced by a mother knowing she has produced one, structured by the class *i* of the supplemental offspring, the class *k* of the mother and the class *j* of the offspring known to have been produced:

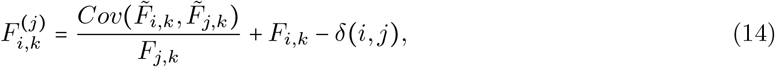

where *δ (i, j)* = 1 if *I = j* and 0 otherwise. This formula (proof in S.M.IV) generalises to a structured generation-overlapping population, the formula for unstructured non-generation overlapping populations of (24) to be found in paragraph 3 of the addendum.

**Same-litter sister matrix** From **P**_**F**_ and eq.14, we can generate **Z**, the same-litter sister matrix, which element *Z*_*i*,*j*_ is the expected number of sisters in class *j* of *ego* in class *i*, by browsing through all classes *k* in which *ego*’s mother may be as she gives birth to this litter: 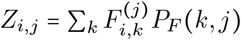, that is:

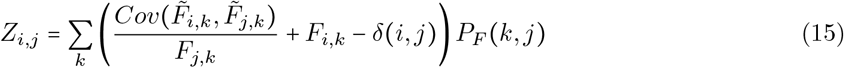

### 3 Substituting the genealogical Markov chain into the kinship formula

Let **D**_*w*_ be the matrix of size, with zeros everywhere but for its diagonal made up of the elements of *w*, the vector of relative abundances. For any *s × s* matrix **A**, (**AD**_*w*_**) (***i, j)* =*A*_*i*,*j*_ *w*_*j*_. Therefore, from eq.13, 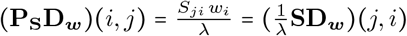 . In other words, 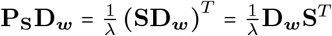, where ^*T*^ denotes transposition. Similarly, we have 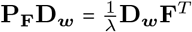 . Now, let 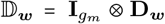 Then, from eqs1,2, classic Kronecker product properties (73) and the above, we get 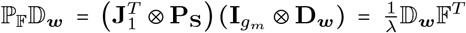 .Similarly we have 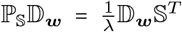 and 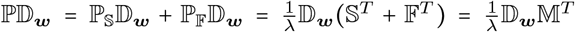 . Let us now multiply both sides of eq.3 by 𝔻_***w***_ (on the right). From the above, we get 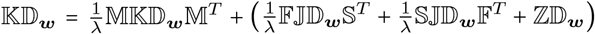 . Note that, in the particular reproductive cases, where ℤ= 𝔽^∗^𝕁ℙ_𝔽_, we have 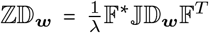 . Call 𝕂_***w***_ = 𝕂𝔻_***w***_, **Z**^†^ = *λ***ZD**_***w***_ and 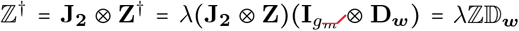 From the above, we get that 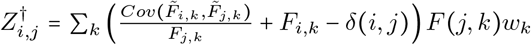 therefore **Z**^†^ can be computed directly from

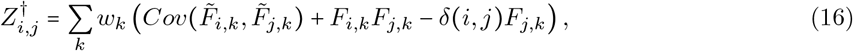

and eq.3 is equivalent to

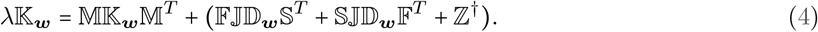

### 4 Vectorization of the kinship formula

An interesting property of the ***vec*** operator and the tensor product is that, for a given set of three matrices with appropriate sizes **A**,**B** and **C**, we have (42; 73):

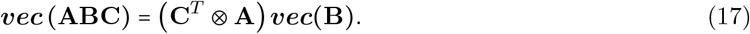

Applied to eq.3, this yields: ***vec*** (𝕂) = (ℙ^*T*^ ⊗ M) ***vec*** (𝕂) + ***vec*** (𝔽𝕁ℙ_𝕊_ + 𝕊𝕁ℙ_𝔽_ + ℤ), that is:

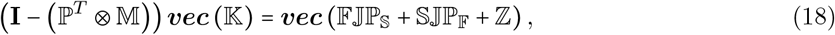

with **I** the identity matrix of size 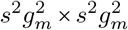 . Vector ***vec*** (𝕂) contains the exact same information than matrix 𝕂, but ordered differently: the *c*_*i+*_*s(g*_*i −1*_)+ (*sg*_*m*_)*(c*_*j −1*_)*+s*^*2*^*g*_*m*_ *(g*_*j*_ −1) element of *vec*(𝕂)of length 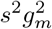, corresponds to the ((*c*_*i*_, *g*_*i*_), (*c*_*j*_, *g*_*j*_)) entry of 𝕂 which can be read in its (*c*_*i*_ + *s*(*g*_*i*_ − 1))^*th*^ row and its (*c*_*j*_ + *s* ( *g*_*j*_ − 1))^*th*^ column and is the expected number of (*g*_*i*_,*g*_*j*_)-kin in class *c*_*j*_ for *ego* in class *c*_*j*_. Applying eq.17 to eq.4, we get: *λ****vec*** (𝕂_***w***_) = (𝕄 ⊗ M) ***vec*** (𝕂_***w***_)+((𝕊 ⊗ F + 𝔽 ⊗ S) ***vec*** (𝕁𝔻_***w***_) + ***vec*** (ℤ^†^))which can be written as

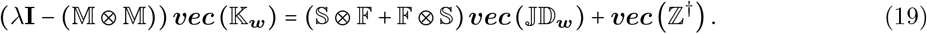

### 5 Existence, unicity and convergence of the kinship formulas

Equation 18 (which is equivalent to eqs.3,4 and 19) admits a solution, and that solution is unique, if and only if (**I** −(ℙ^*T*^ ⊗ M)) is invertible. For any (square) real matrix **A**, its spectral radius (the maximum of the absolute values of its eigenvalues) is such that *ρ*_**A**_<1 if and only if **A**^*t*^ converges exponentially towards **0** as *t* → ∞, its Neumann series (that is,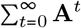 ) converges and (**I** − **A**) is invertible (see 74):

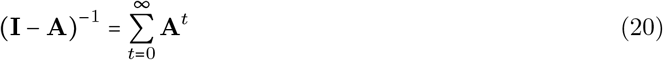

If **S** is a “regular” survival matrix, that is, that it corresponds to the transitions between transient states of an absorbing Markov chain, which absorbing state is “death”, we have lim*t*→+∞ **S**^*t*^ =**0** and *ρ*_**S**_**<**1. Its Neumann series is called the fundamental matrix and is used to compute, for instance, the next-generation matrix (see, e.g., 35). If **S** is a “regular” survival matrix, then 𝕄 also corresponds to the transitions between transient states of an absorbing Markov chain, the absorbing states here, being “death” and generation>*g*_*m*_. This implies that *ρ*_𝕄_< 1. Matrix **P** is the transition matrix of a Markov chain, therefore *ρ*_𝕡_< 1Its generation structured equivalent models the same process, but for individuals with generation number 1, which ancestors via reproduction are not counted, therefore *ρ*_ℙ_< *ρ*_**P**_ =1One spectral property of the kronecker product, is that, for two square matrices of same size **A** and **B**, we have *ρ*_**AB**_ = *ρ*_**A**_*ρ*_**B**_ (75). Therefore *ρ*( ℙ^*T*^ ⊗𝕄) = *ρ*_ℙ_*ρ*_𝕄_ < 1, which completes the proof and we have demonstrated that, for a “proper” survival matrix Eqs4,3 and Eq19 have a (unique) solution, which can be written as:

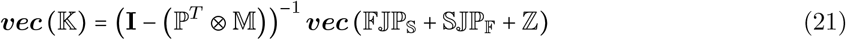

or

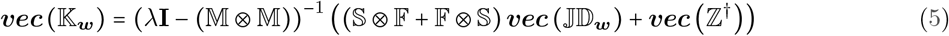

By applying eq.20 and eq.17, to eq.21, we can write this equivalently as an infinite sum which converges:

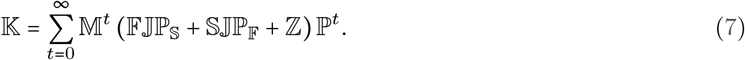

## Supplementary Material

### I Concept of trait

In this manuscript, we call projective traits, or simply *traits*, the categorisation of the *states* of individuals in a structured population. In population ecology, for instance, such traits can be *age, stage*,*body mass, location*,*genotype* etc. In such a population, the population vector is made of several states. The traits categorise these states. In other words, they are the structure of the structure: *traits* are decomposed into *classes* and each individual *state* is a combination of these *classes* . For instance, in a stage×patch model (a 2-trait projection model therefore, also known as a meta-population model following Lebreton (1)), such as the one in (2), each individual *state* can be represented by a pair of *classes*, such as *state* (juvenile, A) which describes the juveniles (individuals in the class “juveniles” of trait stage) in patch A (individuals in class “A” of trait patch).

The concept of projective traits has been rarely used (in population ecology) until recently as most models embedded only one *trait*: such as *age* or *stage*. The concept of multi-trait projection models arises from the increased quantity and quality of data in all fields of population dynamics, which allows an individual in the population to be described by different traits influencing its fate. In (3), the authors develop a theory of such models where they define the “trait structure” of the *r*-trait model as *ts* ={*t*_1_, *t*_2_, .., *t*_*r*_}, where *t*_*i*_ is the number of *classes* of the *i*^*th*^ trait *t*_*i*_. The related projection model is a block-matrix, that can be project over time, like any matrix model (4), the *r*-dimensional population vector, or tensor, providing it is vectorized (5). The result is a *q* × *q* matrix model with *q* = *t*_1_*t*_2_…*t*_*r*_, where the *i*^*th*^ *state* corresponds to the combination of *classes* (*c*_1_, *c*_2_, …, *c*_*r*_) such that 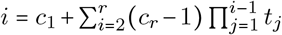 If there are 3 stages (newborn, juvenile and adults) and 2 patches (A and B) in the aforementioned stage × patch model, with *t*_1_=3 and *t*_2_=2, then state (newborn, B)=(*c*_1_=1, *c*_2=_2) corresponds to entry *i=*1+(2 −1)**∗**3= 4 of the population vector of size *q= t*_1_*t*_2_= 6.

Such models allow to analyse the effect of one specific trait, and the process(es) it embeds, on the demography of a population by comparing the model with the trait to “equivalent” models from which the trait is absent. The procedure to obtain the “equivalent” model, called folding by Coste et al. (3), consists in computing, at the demographic stable state, the transition rates between states of the model without the trait under scrutiny by merging all states with the same classes for the other traits. This merger, yielding from projection model **M**, its folded version 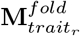, is simply performed by summing all incoming flows and summing all outgoing flows, weighted by the relative abundances of the states to be merged into one.Models 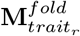 and **M** correspond to the same population: if trait *r* is the colour of the wings of a butterfly, then two entomologists, one colour-blind one not, will obtain, from the exact same data and analysis, respectively models 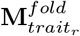 and **M** (6). These have therefore the same asymptotic growth rate and relative abundances but will differ in reproductive values and therefore sensitivities. The analysis of these differences, called Trait-Level Analysis in (3)will describe the effects of trait *r* on the dynamics (and evolution) of the studied population. We illustrate the interest of Trait-Level Analysis when applied to kinship, via the ground squirrel illustration of S.M.V in S.M.V.

### II Discrete time and state space

In many models, such as those of population ecology, the space structure cannot be made continuous (e.g. for a population structured by life history morphs or instars, see 7). At the possible cost of discretisation, the structure consists of *s* classes, and the population can be described by a vector ***n (****t)* of size *s*. Structure discretisation is commonly used in population ecology, when accounting for continuous traits such as body mass, via, for instance, integral projection models (8; 9; 10). For many populations also, the time structure cannot be structured continuously. In population ecology again, for instance, many species, such as most birds or ungulates, reproduce at a specific period (or at specific periods) of the year. In such cases, the models are built from several “seasonal” projectors that are periodic matrices (in the sense of 11) combined (that is, multiplied, if in matrix form) to get an annual (or another time period) projector that does not depend on the year (4).

#### pre- vs post-breeding

If, for instance, reproduction occurs at a specific time-period of the year (modeled by **F’** ), and the rest of the year individual “only” survive (modeled by **S**), one can build an annual model by combining these, either by having reproduction first: **M**_*pre*_ = **S**(**I**_*s*_ + **F**^′^) = **S** + **SF**^′^ = **S** + **F**_*pre*_ or last **M**_*post*_ *=* (**I**_*s*_ *+* **F’) S = S + F’S = S + F**_*post*_. **M**_*pre*_ is called a pre-breeding model and **M**_*post*_ a post- breeding model, they share the same survival (**S**) component but differ in their reproduction components (**F = F**_*pre*_ or **F**_*post*_). As they model the same populations these two models have the same asymptotic properties (same asymptotic growth rate) (see, for instance 12; 4).

### III Independent and identically distributed processes of survival and reproduction

We consider that the process 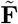 and 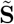 are i.i.d. In other words, we consider that (1) all individuals in class *c*_*i*_ follow the same processes of reproduction 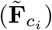 and survival 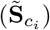, (2) that the individual realisations of survival and fertility are independent from that of other individuals in that same (*c*_*i*_ = *c*_*j*_) or other (*c*_*i*_ ≠ *c*_*j*_) classes and (3) that for a given individual in, the realisation of survival and reproduction are independent: 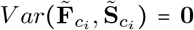 The first assumption of identity and the second assumption of independence are classic in structured population models: the probabilistic fate of an individual is entirely determined by its state and therefore individuals in the same class are considered identical, their fate only differing because of stochasticity. The third assumption, also classic. does not prevent (as it may seem at first sight), to prevent the use of the Kinship Formula for population models that include, for instance, survival/fertility trade-offs. However, it can be easily shown that any model can be made to obey assumption (3), by adapting its structure (13). We illustrate it here for two cases that are common in population ecology.

Consider the case of “seasonal” matrices described in S.M.II. As is obvious from the formulas of **F = F**_*pre*_ *=* **SF’** and **F = F**_*post*_ *=* **F’S**, the reproductive process at the annual level **F** depends, in general, on the survival process. However, while the post-breeding reproduction, at the annual level, via 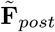, of an individual in class *c*_*j*_ depends on the realisation of 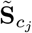 for that same individual, it is not the case for the pre-breeding reproduction. 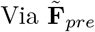, the first seasonal stochastic matrix encountered is 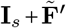, where the first component is constant and the second one yields different individuals by definition. Therefore, in the case of seasonal models, using pre-breeding formulation, allows to respects the third assumption. On a side note, the extension of the kinship formula to account for varying environments (in prep), and in particular to compute the kinship of a population modelled via periodic matrices (in the sense of 11), such as **M**_*post*_ *=(***I**_*s*_ **F’)S** will only require independence of survival and fertility at the level of each periodic matrix. Here survival and fertility are clearly independent for both **S** and (**I**_*s*_ **F’)** as each contain only one such process, and therefore the kinship of the population modelled by **M**_*post*_ will be possible and provide the kinship both after the fertility event and after the survival event.

The second case seemingly preventing assumption (3) is the case of trade-offs between *realisations* of survival and reproduction. In (14), such trade-offs are called “individual” trade-offs (which differ from “population” trade-offs which occur between *expectations* of survival and reproduction, across classes). Those can be modelled by having, for instance, for an individual in class *c*_*j*_ in a given year, survival 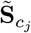 that year be a function of reproduction 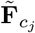 that year, which violates the third assumption above. Here again, a careful choice of the structure allows to circumvent the problem, by adding trait *realised reproduction* to the structure. This is commonly done in bird population ecology, where, by adding classes “breeders” and “non-breeders” in the structure, one segregates individuals producing offspring or not during the current year. The trade-off is then incorporated by having “breeders” and “non-breeders” have different survival processes. In such a model, the stochasticity in the realisation of fertility – will I breed or not this year? – is therefore embedded in the survival of the previous time-step –will I breed or not next year ? that is, this current year, in case I survive, am I transitioning towards the “breeders” or the “non-breeders” class ? (in a structured model, survival involves both survival proper, and the transition of survivors to the various stages). In that case, there is independence, for a given individual in class *c*_*j*_ at a given time-step, between the realisation of its fertility (which can be constant, or depend on offspring survival or breeding success, but does not influence survival) and its survival, and assumption (3) is respected (see 15, for an example of such a model).

### IV Same Litter Fertility

In order to count the number of kin, we count the descendants of the ancestors of *ego*. We can reach the descendants stemming from the older and younger sisters of the daughter of the ancestor by respectively. applying to the ancestor one backward survival projection (via **P**_**S**_ or ℙ_𝕤_ to consider the ancestor *before* it produced its daughter in the genealogy of *ego*) followed by general forward projection (via **M** or 𝕄) or one forward survival projection (via **S** or 𝕤 to consider the ancestor *after* it produced its daughter in the genealogy of *ego*) followed by general forward projection. In other words, the expected number of kin via older and younger sisters of the ancestor’s daughter only depend on the deterministic projection model **M= F+S**.

However when counting the descendants via sisters of the ancestor’s daughter, it proves necessary, in the general case, to consider a further building-block, the same-litter-sister matrix **Z**, which element *Z*_*i*,*j*_ is the number of sisters of the same-litter in class *j* of an individual in class *i* that has just been born. **Z** can be inferred from 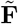 via the following steps:

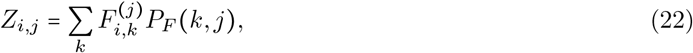

where 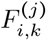 is the number of same-litter daughters, that is, the expected number of supplemental offspring produced by a mother knowing she has produced one, structured by the class *i* of the supplemental offspring, the class *k* of the mother and the class *j* of the offspring known to have been produced. Equation 22 simply equates the number of same-litter sisters of *ego* in class *j* to the number of same-litter daughters of the mother of *ego* weighted by the probability distribution of the class of the latter. The number of same-litter daughters depends, from its definition, on 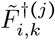 which has the distribution of 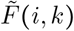 biased by 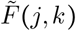 and on whether *ego* and the kin are in the same class:

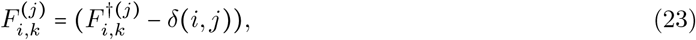

where *δ*(*i, j*) = 1 if *i* = *j* and 0 otherwise, and where

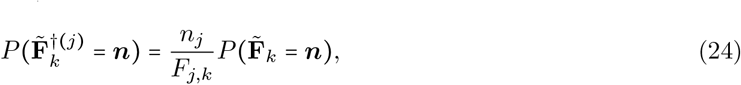

where for a matrix **A, A**_*k*_ is the vector corresponding to the *k*^*th*^ column of **A**, that is to {*A*_*i*,*k*_}_*i*_.

From this we can compute the expectation,

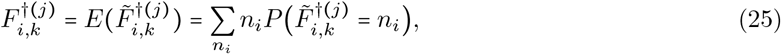

by noting that 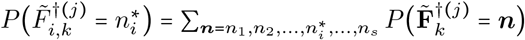 From eq. 24, we get therefore

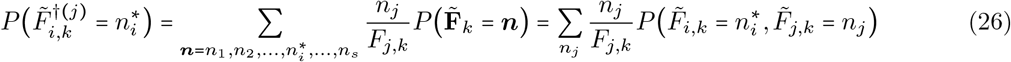

From Eqs 26 and 25, we get therefore

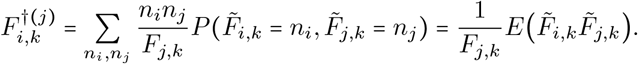

That is,

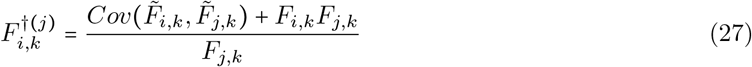

From Eqs 27 and 23, we then have

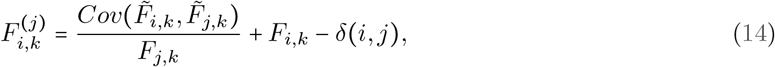

Note that this formula differs from and corrects Eq(9) of (16) which was erroneous despite the calculations being “elementary”. From Eqs 14 and 22, we get finally

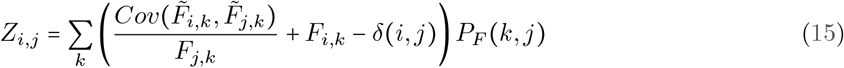

#### Particular reproductive cases

In three particular (but very common in the literature) cases, the number of same-litter daughters does not depend on the class of the daughter ancestor of *ego*: 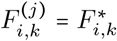 The same-litter daughters is now 2-dimensional and can be represented as a matrix 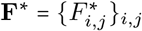 eq. 22 becomes 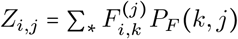 ,which can be written in matrix form as **Z=F**^**∗**^**P**_**F**_, which relates these particular common cases with the general case. These three cases are:

#### Bernoulli Reproduction

In that case, individuals can produce at most 1 offspring per time step: there is by definition no same-litter daughters, then we simply have

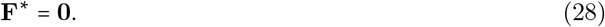

#### Independent Poissons reproduction

Here 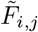 is a Poisson independent from 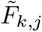, therefore 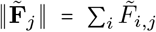 is a Poisson as the sum of *s* independent Poisson. A known result is that, in that framework, the number and state of supplemental offspring is independent from the number and states of offspring produced and therefore

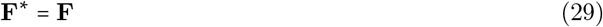

This can be checked directly. From Eq14 we have, 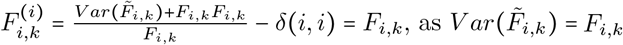 and *δ*(*i, i*) = 1. For *i* ≠ *j*, the same equation gives 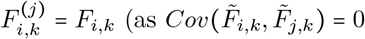 and *δ*(*i, j*) = 0) which is indeed independent from *j*. Therefore, in all cases, 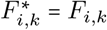

#### One offspring class

If there is only one class of offspring (say class 1), then **F**^**∗**^ is non-zero only for 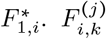 is indeed only non-zero for *i* = 1 and *j* = 1, we have 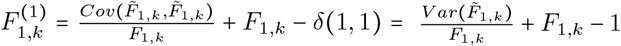 . Therefore **F**^**∗**^ is such that

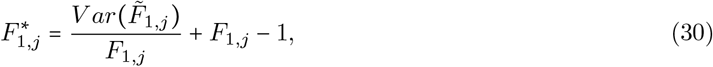

and 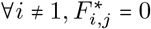

### V Illustration

Here we illustrate our method on a population of ground squirrels *Spermophilus dauricus*, structured by stages. It is an extended Leslie model, i.e. an age-structured model with a class that regroups all individuals older than a certain age. All the computations can be reproduced from the main text equations or directly from the implementation of the equations of the main text (in R and in Matlab) to be found in S.M.VIII.

Here, classes 1 and 2 correspond to individuals that are about to turn 1 and 2 year(s) old, respectively, and class 3 corresponds to older individuals. We want to compute the expected number of all (*g*_*i*_ − *g*_*j*_)- kin for all *g*_*j*_ and *g*_*i*_ ≤ *g*_*m*_ = 3; there are 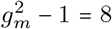 such kin: the mother (1,2), grand-mother (1,3), the daughters (2,1), grand-daughters (3,1), the sisters (2,2), maternal aunts (2,3), nieces (3,2) and cousins (3,3) (see fig.1). From the life table for that population from (17), we get the following projection matrix:

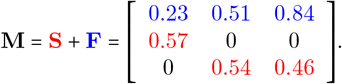

#### Generation-structured matrices

From these and via eqs 1 and 2,we can construct 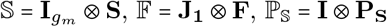 and 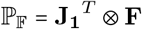 for *g*_*m*_ = 3. That is, 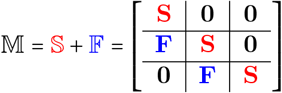 (see below for ℙ_𝕊_ and ℙ_𝔽_).

#### Kinship Matrix

Without further information, we assume that the fertility is Poisson and therefore **F**\* = **F** and 𝔽* = 𝔽. Via a simple eigen-analysis, we obtain the asymptotic growth rate *λ* = 1.00 (we round all numbers to 2 digits, note however that this asymptotic growth rate is < 1) and the stable class distribution **w** = (0.47, 0.27, 0.27). From the latter, we can construct **D**_***w***_ the matrix with zeros everywhere but for its diagonal made up of the elements of ***w***, and from it 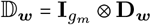. We now have all the elements to generate, from eq. 8, 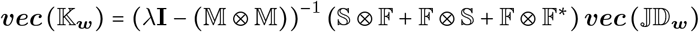, where 𝕁 = **J**_**3**_⊗ **I**_***S***_, with **J**_**3**_, the zero *g*_*m*_ × *g*_*m*_ zero-matrix with 1 in the (1,1) position). From the solution to this equation we easily get (see main text for interpretation)

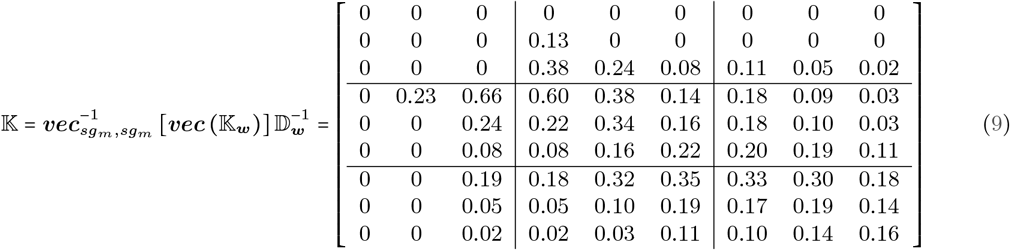

#### Unstructured kinship

From equation 10, we get the unstructured kinship matrix, the symetric matrix which gives the expected number of any kin for *ego* taken at random in the stable state population:

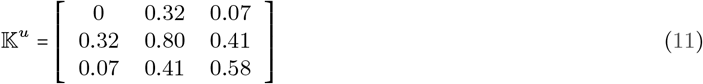

Reading **K**_**u**_, we know that *ego* is expected to have 0.80 sisters, 0.58 cousins, 0.41 nieces and aunts and 0.07 grand-daughters and grandmothers (that is, the probability that its grandmother is alive is 0.07)

#### Kinship of unstructured population

In order to illustrate one of the uses of these calculations in ecology and evolution, we generate **M**^*u*^ = **F**^*u*^ + **S**^*u*^ the unstructured model equivalent to **M = F + S** (see S.M.I). **M**^*u*^ is a scalar, that is a 1 × 1 matrix which says that all individuals in the population have the same probability to survive **S**^*u*^ = 0.53 and the same expected number of offspring **F**^*u*^ = 0.47. This model has the same asymptotic growth rate than **M**: *λ* = **M**^*u*^ = 1.00, as they model the same population. However, the kinship matrix of **M**^*u*^, generated assuming similarly Poisson reproduction yields a different output:

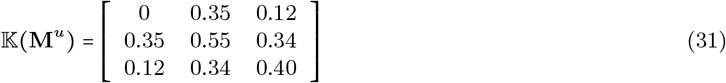

By comparing the kinship matrix of the unstructured population (eq.31) and the unstructured kinship of the structured population (eq.11), we can draw conclusions about the effect of the processes embedded in the classes of the model on kinship. Here the classes correspond to age classes (broadly speaking, the last one being open ended and therefore a “stage”), and they embed mostly, an increase in fertility with age and a decline in survival with age (senescence). The combination of these effects implies that the probabilities that direct ancestors (mother, grandmother) will be alive and the expected number of direct descendants (daughters, grand-daughters) will be lower than for the equivalent model without individual heterogeneity, where all individuals have the same vital rates (S.M.I). To the contrary, it implies that the number of other, indirect, kin, the descendants of ancestors (sisters, nieces, cousins and aunts) are more numerous. In other words, there is a trade-off between the number of direct and indirect kin as a response at the kinship level of the trade-off embedded in the life history of the population (increased allocation with age towards reproduction at the cost of survival).

#### Genealogical Markov chains

These are intermediate tools that are not necessary for the computation of the Kinship Matrix via eq 8 or eq 5. However they appear in the Kinship Formula itself (eq 3), we therefore provide their expression for the illustration, here. From *λ* and **w**, we can construct **P** from eq13, 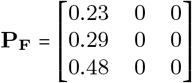 and 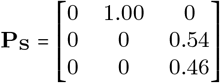. From these, it is easy to build 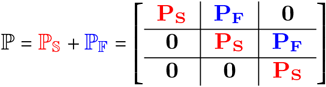

### VI The kinship formula as an infinite sum

Eq.3 can also be written as

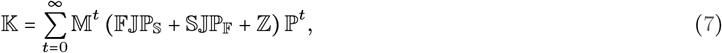

This can be seen by by applying eq.20 and eq.17, to eq.21, or, and it is easier, by multiplying both sides of eq.7 on the left by 𝕄 and on the right by ℙ and via the *T* = *t* + 1 change of index. The sum in eq.7 converges iff **S** does. This formulation as an infinite sum is the generation structured equivalent of that in (16) and corresponds to the natural approach to Kinship Demography when one considers the ancestor of *ego* at all time-steps in the past (18; 19). Contrary to these approaches however,eq.7 accounts for all kin relationships (up to the arbitrary distance *g*_*m*_) simultaneously. Matrix ℙ^*t*^ provides the probability distribution of the *g*_*j*_ ancestor of *ego t* time-steps before present. The term in parenthesis then considers younger sisters, older sisters and same-litter sisters of the *g*_*j*_ − 1 ancestor, and 𝕄^*t*^ accounts for their descendants today. Kinship Matrix 𝕂 differs from and extends the matrix of the same name in (16), where it corresponds to a specific kin relationship. Equation 7, like eq. 5, and contrary to eq.3, provides a direct expression for 𝕂.

#### Embedded maximum age

Consider the case where there is a maximum age embedded in the model (i.e., it is age-structured), that is there exist *ω* whereby **S**^*ω*^ = **0** (but **S**^*ω −*1/^ ≠ **0**).In that case 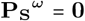 as well (if one cannot live more than *ω* time-steps, one cannot have been alive more than *ω* time-steps ago). This is also true in the generation structured model: 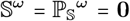 To the contrary, **F** and 𝔽 do not behave similarly, and contrary to **F**, we have for the generation structured model 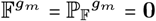 Any product of a series of 𝔽 and 𝕊 such as 𝕊𝕊𝔽𝔽𝔽𝕊𝔽, of any size, that contains more than *g*_*m*_ 𝔽s is worth **0** since 𝔽 increases the generation number by 1 and 𝕊 preserves it. Therefore, as it contains either a run of 𝕊s longer than *ω* −1 or more than *g*_*m*_ −1 𝔽s, we have 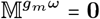 and similarly 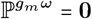 . Each element of (𝔽𝕁ℙ_𝕊_ + S𝕁ℙ_𝔽_ + 𝔽^**∗**^𝕁ℙ_𝔽_) contains either one 𝔽or one ℙ_𝔽_ and one 𝕊 or one ℙ_𝕊_ and therefore

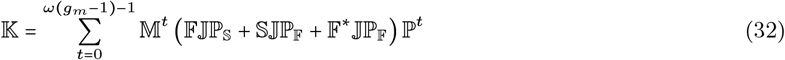

### VII. Eventual number of kin

The Kinship Formula, as formulated in eq 7, provides the *current* number of kin of *ego*, that is, the number of kin that are currenlty alive. It may be interesting to know also the *eventual* number of kin, that is, the total number of kin that *ego* will eventually have, including those which died before *ego* was born and those who will be born after *ego*’s death. To achieve this, one natural first step is to consider the non-generation-overlapping version of the projection model **M=F+S**. It corresponds to the next-generation matrix **R=F(I −S)**^**−**1^ (4). For simplicity, let us consider the case where there is only one class of offspring. Then, over a generation, we have **F** = *R*_0_, a scalar, the net reproductive rate, corresponding the (1,1)entry of **R**. We also have simply that **S =** 0. From these, we get **P**_**F**_=1, **P**_**S**_=0 and *ω=*1. We are in the case here with only one offspring class (as the model has only one class) and therefore we can compute the number of same-litter daughters/sisters (these are the same here as **P**_**F**_ = 1) which is a scalar: 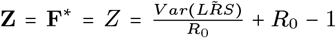, where 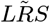 is the Lifetime Reproductive Success, the expectation of which is *R*_0_ (this can be computed from 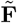 and **S** via, for instance, Markov chains with rewards (see e.g. 20)). From eq 32, we can compute the Kinship Matrix for this model: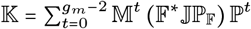, which yields a diagonal matrix such that 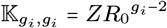 and 𝕂_1,1_ =0 In other words, *ego* is expected to have 𝕂_2,2_ =*Z* sister, 𝕂_3,3_=*ZR*_*0*_ (first cousins),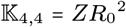 etcs. By construction, apart from these kin of the same generation than *ego*, and therefore on the diagonal of 𝕂, there is no other kin alive as generations do not overlap: *ego*’s mother and aunts are dead already and its daughters and nieces are not born yet.

#### eventual number of kin in the general case

To remedy this, and be able to compute the eventual number of kin, we add class “death” to the model. Obviously, in this case, the “survival” matrix of the model includes dead individual as well, and therefore **S**^*t*^ does not converge as *t*→ ∞. In the general case, this could imply that the kinship formula does not converge, however, we are here in a particular case where the individuals that are allowed to live forever cannot reproduce (that is, here, individuals in the class “death”). The kinship formula then converges and the kinship matrix is made easier to read by ensuring that all individuals to be counted are in the “death” class. This is done by summing the summands of eq.7 from *t* = 0 up to *t = ω* (*g*_*m*_ − 1)–1 + *ω*) (instead of *t=ω (g*_*m*_ −1) – 1 as per eq32 ). Let us illustrate this for our non-generation overlapping, unstructured example.

#### illustration: eventual number of kin in a non-generation-overlapping population with one class of offspring

In this case, the components of the projection matrix become 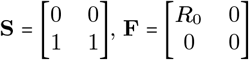 and 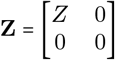 Similarly, we have 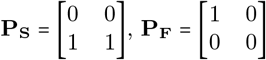. And remembering that here, *ω* = 1, we get

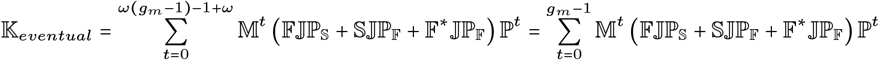

which, for *g*_*m*_ = 4 yields

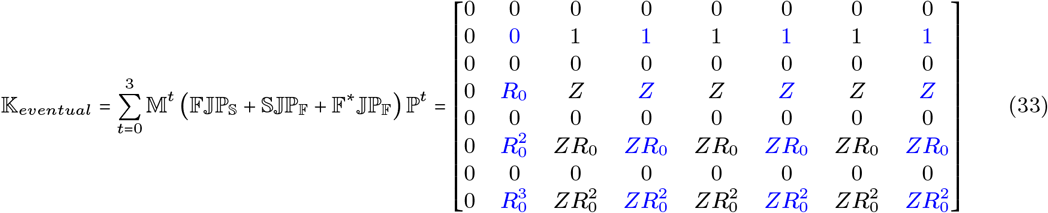

In this kinship matrix, we are only interested in the “death” class for both the kin and *ego* (those are highlighted in blue in eq 33), that is:

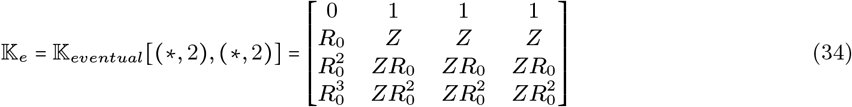

### VIII Code

**R Code**

~~~
# Inputs
F = matrix(c(0.23, 0.51, 0.84, 0, 0, 0, 0 ,0 ,0),nrow=3,byrow=T) #fertility component of matrix
S = matrix(c(0, 0, 0, .57 ,0 ,0,0, .54 ,.46),nrow=3,byrow=T) #survival component of matrix gmax=3 #number of generations investigated
#Z= ? #same-litter-sister matrix
# eigen properties of the population projection matrix M=F+S #projection matrix
ev <- eigen(M);lam <- max(Re(ev$values));
w_raw <- abs(Re(ev$vector))[, which.max(Re(ev$values))] w <- w_raw/sum(w_raw)
#S<-myDiag(matrix(0, omega, omega), rep(1, omega-1), -1) %x% S # genealogical matrices
# w[w<0]<-0
#P=t((1/lam)*A*(w^(-1)%*%t(w))) #P[is.nan(P)]<-0
#PF= (t(F)/t(A))*P
#PF[is.nan(PF)]<-0 #PS=P-PF
# Matrix Z
#POISSON reproduction Fstar=F
#BERNOULLI reproduction #Fstar=matrix(0, dim(F)[1],dim(F)[1])
#1 CLASS OF OFFSPRING
#Fstar=
#Z=Fstar%*%PF
# Generation structured matrices #s=dim(F)[1]
myDiag <- function(x, vec, k) { x[row(x) == col(x) - k] <- vec
 x } #turns a vector (vec) into a zero matrix matrix but with vec on the kth diagonal J1<-myDiag(matrix(0, gmax, gmax), rep(1, gmax-1), -1)
bigF=J1%x%F
bigS=diag(gmax)%x%S bigM=bigF+bigS
bigFstar=J1%x%Fstar
#temp=matrix(0, gmax, gmax);temp[2,2]=1; #bigZ=temp%x%Z
# kinship matrix
temp=matrix(0, gmax, gmax);temp[1,1]=1;s=dim(F)[1] J=temp%x%diag(s)
Dw <- diag(w)
bigDw <- diag(gmax)%x% Dw
vec <- function(x) { y <- as.vector(x)
 return(y) }
vecKw=solve(diag(s*s*gmax*gmax)-bigM%x%bigM)%*%(bigS%x% bigF + bigF%x% bigS+ bigF%x% bigFstar)%*%vec(J*bigDw) Kw <- matrix(vecKw, nrow = gmax * s, ncol = gmax * s)
K <- matrix(vecKw, nrow = gmax * s, ncol = gmax * s) %*% solve(bigDw)
Jy <- diag(gmax)%x%matrix(1, nrow = 1, ncol = s)
Ku <- matrix((Jy%x%Jy)%*%vecKw, nrow = gmax, ncol = gmax)
~~~

**Matlab Code**

~~~
%% inputs
%components of the projection matrix
F =[ 0.2300 0.5100 0.8400; 0 0 0; 0 0 0]
S = [ 0 0 0; 0.5700 0 0; 0 0.5400 0.4600]
%kinship distance investigated gmax=3
%% Basic eigen-analysis M=F+S %projection matrix
[w,lam]=eigs(M,1); %eigen analysis lam % asymptotic growth rate
w=w/sum(w) %asymptotic relative abundances
%same-litter daughters Fstar=F;
%% Generation structured matrices/tensors ss=size(F,1);
J1=diag(ones(1,gmax-1),-1);
bigF=kron(J1,F); bigS=kron(eye(gmax),S); bigM=bigF+bigS
bigFstar=kron(J1,Fstar);
temp=zeros(gmax,gmax);temp(1)=1;ss=length(F); J=kron(temp,eye(ss));
Jy=kron(eye(gmax),ones(1,ss));
%% Direct Computation of expectation (of kinship matrix) Dw=diag(w);
bigDw=kron(eye(gmax),Dw);
vecKw=((lam*eye(ss*ss*gmax*gmax)- kron(bigM,bigM))^(-1)) * (kron(bigS,bigF)+kron(bigF,bigS)+kron(bigF,bigFstar))*vec(J*bigDw);
Kw=reshape(vecKw,[gmax*ss,gmax*ss]);
K=reshape(vecKw,[gmax*ss,gmax*ss])*(bigDw^(-1)); Ku=reshape(kron(Jy,Jy)*vecKw,[gmax,gmax]);
format shortg round(K,2) round(Ku,2)
%% vec operator
function [y] = vec(x)
  y=x(:);
end
~~~

